# A Seed-Derived Multi-Kingdom Synthetic Community Enhances Radish Seedling Protection Against *Alternaria brassicicola* Abra43

**DOI:** 10.64898/2026.10.01.755899

**Authors:** Natalia Guschinskaya, Amandine Le Ruyet, Justine Colou, Gontran Arnault, Isabel Guerrero Zepeda, Coralie Marais, Muriel Marchi, Logan Suteau, Philippe Simoneau, Claire Campion, Marie Simonin

**Affiliations:** Univ Angers, Institut Agro, INRAE, IRHS, SFR QUASAV, F-49000 Angers, France

## Abstract

Seed-associated microorganisms represent a promising resource for developing microbiome-based strategies against seed-borne pathogens. Here, we assembled synthetic microbial communities (SynComs) from cultivable members of the natural radish seed microbiota and evaluated their capacity to protect seedlings against the seed-transmitted fungal pathogen *Alternaria brassicicola* Abra43. Seven SynComs representing single-or multi-kingdom combinations of bacteria (B), filamentous fungi (F), and yeasts (Y) (B, F, Y, BF, BY, FY, and BFY) were inoculated onto radish seeds. We assessed their effects on disease incidence, seedling growth, microbial community assembly (ITS1 and *gyrB* gene metabarcoding) and pathogen colonization (metabarcoding and specific qPCR on *dhn1* gene for *A. brassicicola* Abra43).

Protection of radish seedlings against *A. brassicicola* Abra43 strongly depended on SynCom composition. The yeast fraction alone provided little protection, whereas the bacterial fraction partially reduced symptoms and maintained seedling growth despite substantial pathogen colonization, suggesting increased plant tolerance. SynComs containing filamentous fungi markedly reduced *A. brassicicola* Abra43 abundance, and the full multi-kingdom BFY SynCom provided the most consistent protection, combining reduced disease incidence, restored seedling growth, and low pathogen relative and absolute abundance. Inoculated strains, *Alternaria alternata* ABAI001 and *Plectosphaerella* sp. FL19 were enriched in protective SynComs and displayed antagonistic activity against *A. brassicicola* Abra43 *in vitro*. Together, these results show that seed-derived multi-kingdom communities can reshape early plant microbiota and that combining bacterial and fungal strains with complementary ecological functions can improve protection against a seed-borne fungal pathogen.

---

Seeds are a critical interface between plant propagation, microbiota transmission, and pathogen spread. In addition to carrying the plant embryo, seeds harbor microbial communities that can contribute to early plant establishment, seedling vigor, and protection against disease (Matsumoto et al. 2021; Colaert-Sentenac et al. 2026). These seed-associated microorganisms may be transmitted to emerging seedlings, but germination imposes strong ecological filtering through rehydration, tissue expansion, and rapid changes in nutrient availability (Barret et al. 2015; Olofintila and Noel 2023; Gerna et al. 2026). Consequently, only part of the seed microbiota is expected to persist during the seed-to-seedling transition (Rochefort et al. 2021; Walsh et al. 2021), and this early selection may determine whether beneficial microorganisms establish in time to interfere with seed-borne pathogens.

Seed-borne diseases are particularly difficult to control because infection can begin before and during germination, when emerging seedlings are highly vulnerable and their specific immune responses may favor susceptibility to necrotrophic pathogens (Ortega-Cuadros et al. 2024). *Alternaria brassicicola* Abra43 (hereafter Abra43), a necrotrophic fungal pathogen of *Brassicaceae* crops, causes black spot disease and damping-off and can reduce seed quality, germination, seedling growth, and crop establishment (Chirco and Harman, 1978; Pattanamahakul and Strange, 1999; Pochon et al., 2012; Guillemette et al., 2014). Infected seeds represent an important primary inoculum source, allowing the pathogen to persist between cropping cycles and initiate disease at early developmental stages. Because resistant *Brassicaceae* cultivars remain limited and pesticide reduction is an important agricultural objective, alternative approaches are needed to protect seedlings during this early infection window (Iacomi-Vasilescu et al., 2004; Saharan et al., 2016; de Britto and Jogaiah, 2022; Kaur and Dutta, 2024a, 2024b).

The natural seed microbiota represents a potential source of protective microorganisms (Nelson, 2018; White et al., 2018; Barret et al., 2019). Seed-associated bacteria have already been linked to resistance against seed-borne pathogens, as shown for *Sphingomonas melonis*, which protects rice against *Burkholderia plantarii* (Matsumoto et al., 2021). In *Brassicaceae*, seed bacterial community structure can also influence interactions between beneficial microorganisms and pathogens (Rybakova et al., 2017; Walitang et al., 2017). Synthetic microbial communities, or SynComs, offer a way to test selected members of this microbiota under controlled conditions and to identify microbial combinations that contribute to plant protection. However, most protective SynCom studies have focused on bacteria (Xu et al., 2025), whereas filamentous fungi and yeasts may also contribute to pathogen suppression especially against fungal pathogens, and overall plant health.

Multi-kingdom interactions may be particularly important for seed microbiota function. Seed microbiota are characterized by the close coexistence of bacterial and fungal communities, which can display comparable levels of diversity (Simonin et al., 2022). This cross-kingdom coexistence creates frequent opportunities for interactions among bacteria, filamentous fungi, and yeasts, although their functional significance remains poorly understood. Previous studies have shown that bacteria can influence fungal and oomycete communities in plant-associated microbiota (Durán et al., 2018), and that cross-kingdom bacterial-fungal SynComs can enhance the suppression of fungal diseases such as Fusarium wilt when directly inoculated onto germ-free tomato seedlings (Zhou et al., 2022). Related work has also demonstrated the disease-suppressive potential of reconstructed rhizosphere communities against Fusarium wilt (Liu et al., 2026). However, these studies primarily focused on established seedlings or rhizosphere communities rather than on microbial persistence across the seed-to-seedling transition. This distinction is particularly relevant for seed-applied SynComs, because inoculated microorganisms must survive seed inoculation, persist during germination, colonize emerging seedlings, and retain their protective functions while the seed-borne pathogen becomes active. Although seed-to-seedling transmission is strongly filtered and taxon dependent (Rochefort et al., 2021; Chesneau et al., 2022), whether microorganisms originating from the seed microbiota are particularly competent at crossing this ecological transition remains poorly understood.

Here, we assembled SynComs from cultivable bacteria, filamentous fungi, and yeasts isolated from the natural microbiota of radish seeds and evaluated their ability to protect seedlings against the seed-borne pathogen *A. brassicicola* Abra43. The SynComs comprised eleven bacterial, six filamentous fungal, and five yeast strains and were applied directly to radish seeds as single-group communities (B, F, or Y), two-group combinations (BF, BY, or FY), or a complete three-group consortium (BFY), in the presence or absence of Abra43.

We tested whether these SynComs could establish in seeds and seedlings, reshape bacterial and fungal community composition, reduce Abra43 colonization, limit disease symptoms, and preserve seedling growth. By combining phenotyping, metabarcoding-based community monitoring, and pathogen-specific qPCR, this study evaluates the potential of seed-derived multi-kingdom SynComs for biological protection against a seed-borne fungal pathogen.

We first assessed whether seed-applied SynComs could protect radish seedlings against *Abra43* by combining disease scoring, seedling height measurements, ITS1-based estimation of Abra43 relative abundance, and absolute qPCR quantification of the pathogen-specific *dhn1* marker (Fig. 1). In the Abra43-inoculated water control (H^2^O +), disease was severe, with 95 ± 5% symptomatic seedlings (Fig. 1A), and seedling height was strongly reduced compared with the non-inoculated control (1.88 ± 0.63 versus 3.45 ± 0.66 cm; Fig. 1B). Abra43 also dominated the fungal community, accounting for 99.38 ± 0.23% of ITS1 reads (Fig. 1C), and was detected at high abundance by pathogen-specific qPCR (Fig. 1D). Together, these results confirmed efficient pathogen establishment and its strong impact during radish seedling emergence.

**Fig. 1.**
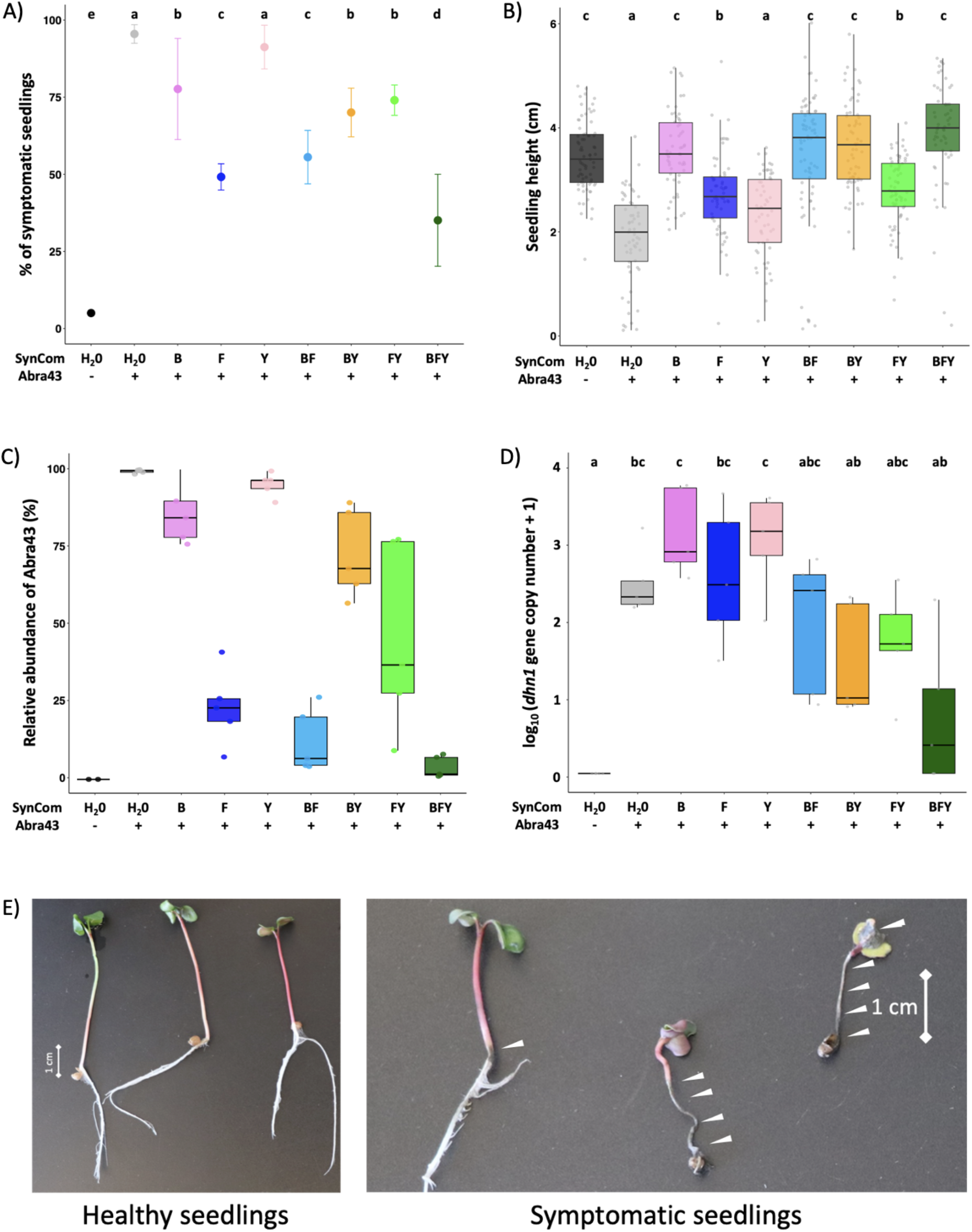
Effect of SynCom treatments on: A) Percentage of symptomatic seedlings. Bars represent the mean ± standard error (SE). B) Seedling height. Boxplots represent the distribution of seedling height from the collar at the base of the cotyledons (cm) (n=100 seedlings). C) Relative abundance of Abra43 in the fungal community, determined by ITS1 region metabarcoding and expressed as the relative abundance of *A. brassicicola* ASV. (D) Absolute abundance of Abra43 quantified by qPCR using the pathogen-specific *dhn1* marker and expressed as log^10^ (*dhn1* gene copy number + 1). The central line indicates the median; boxes represent the interquartile range; whiskers show data variability; and gray points correspond to individual biological replicates (n=5). For A, B, C and D panels, different lowercase letters above boxes indicate significant differences according to the Dunn post hoc test (Benjamini-Hochberg correction, *p* < 0.05). Treatments are identified according to their SynCom composition: H2O (water control), B (bacteria), F (filamentous fungi), Y (yeast), BF (bacteria + filamentous fungi), BY (bacteria + yeast), FY (filamentous fungi + yeast), and BFY (bacteria + filamentous fungi + yeast). The symbols “-” and “+” below the x-axis indicate the absence (-) or presence (+) of *Abra43*. E) Pictures of representative healthy and symptomatic radish seedlings. The arrowheads indicate necrotic lesions specifically associated with Abra43.

SynCom-mediated protection strongly depended on community composition. The yeast-only SynCom showed no evidence of protection, with 89 ± 12% symptomatic seedlings, reduced seedling height (2.33 ± 0.78 cm), and high Abra43 relative abundance (95.14 ± 1.69%). In contrast, the bacterial and filamentous fungal fractions produced statistically distinct protective phenotypes. The bacterial SynCom partially reduced symptom incidence and maintained seedling height close to that of the non-inoculated control (3.56 ± 0.74 cm), despite a high Abra43 relative abundance (85.64 ± 4.34%). This uncoupling between pathogen colonization and plant performance is consistent with a predominantly tolerance-like effect of the bacterial fraction rather than strong restriction of pathogen establishment. Similar growth-maintaining effects have been reported for plant-associated bacterial communities: bacterial commensals can modulate phytohormone homeostasis and preserve root development (Finkel et al., 2020), or alleviate immune-associated growth inhibition by modulating the plant growth-defense balance (Ma et al., 2021). Thus, the near-control growth observed here despite substantial Abra43 colonization may reflect bacterial modulation of host physiology rather than direct pathogen suppression.

By contrast, the filamentous fungal SynCom was associated with a stronger reduction in symptom incidence (54.86 ± 15.83%) and a marked decrease in Abra43 relative abundance (23.23 ± 5.48%), although seedling growth was only partially restored (2.72 ± 0.84 cm). This suppression may reflect direct fungal-fungal interactions, including competition for space and nutrients, niche pre-emption, antibiosis, or mycoparasitism, all of which have been described as mechanisms underlying fungal biocontrol of phytopathogens (Köhl et al., 2019; Rajani et al., 2021). Interestingly, seed-associated filamentous fungi isolated from wild *Brassicaceae*, including Alternaria spp., have also been shown to antagonize *A. brassicicola* Abra43 *in vitro* (Lerenard et al., 2024), supporting the potential contribution of resident seed-associated fungi to pathogen suppression.

All two-group SynComs provided partial phenotypic protection compared with the Abra43-only treatment, but none fully restored the phenotype of non-inoculated plants. However, their effects differed more markedly at the level of pathogen colonization, with BF strongly reducing Abra43 relative abundance to 12.44 ± 4.58%, despite only partial restoration of seedling growth (3.48 ± 1.23 cm). In contrast, BY and FY were still associated with high symptom incidence (69 ± 17% and 76 ± 12.55%, respectively), highlighting that phenotypic and molecular indicators of disease suppression were not fully concordant across SynComs.

The complete three-groups BFY SynCom provided the most consistent protective phenotype, combining the lowest proportion of symptomatic seedlings among Abra43-inoculated treatments (35 ± 2.5%), restored seedling height comparable to the non Abra43-inoculated condition (3.89 ± 1.04 cm), low Abra43 relative abundance (3.86 ± 1.53%), and the lowest observed pathogen signal by qPCR. These results indicate that protection depends not only on the activity of individual microbial fractions but also on their combination, consistent with complementary contributions of bacteria, filamentous fungi and yeasts. Similar functional complementarity has been reported in tomato, where combined bacterial-fungal SynComs provided stronger suppression of Fusarium wilt than bacterial or fungal communities alone (Zhou et al., 2022). More generally, microbial communities with greater diversity can be more resistant to invasion, potentially because broader resource use and niche occupation reduce opportunities for pathogen establishment (van Elsas et al., 2012; Mallon et al., 2015). However, invasion resistance also depends strongly on community composition and interaction structure rather than diversity *per se*, as shown for plant-associated microbial communities challenged with *Ralstonia solanacearum* (Wei et al., 2015; Li et al., 2019).

Metababarcoding analysis revealed that fungal community composition was strongly structured by both SynCom treatment and habitat (seed *vs* seedling, Fig. 2A). Principal Coordinates Analysis (PCoA) based on Bray-Curtis dissimilarities of ITS1 profiles revealed significant effects of SynCom composition (PERMANOVA, R^2^=0.38, p=0.0001) and habitat (seed *versus* seedling; R^2^=0.18, p=0.0001). The larger proportion of variation explained by SynCom composition indicates that fungal communities retained a strong signature of the inoculated community during the seed-to-seedling transition. Taxonomic profiles nevertheless revealed substantial restructuring after germination (Fig. 2B). Abra43 remained highly dominant in H2O, B, and Y treatments inoculated with pathogen, and, to a lesser extent, in BY, whereas treatments containing the filamentous fungal fraction were associated with markedly lower pathogen dominance.

**Fig. 2.**
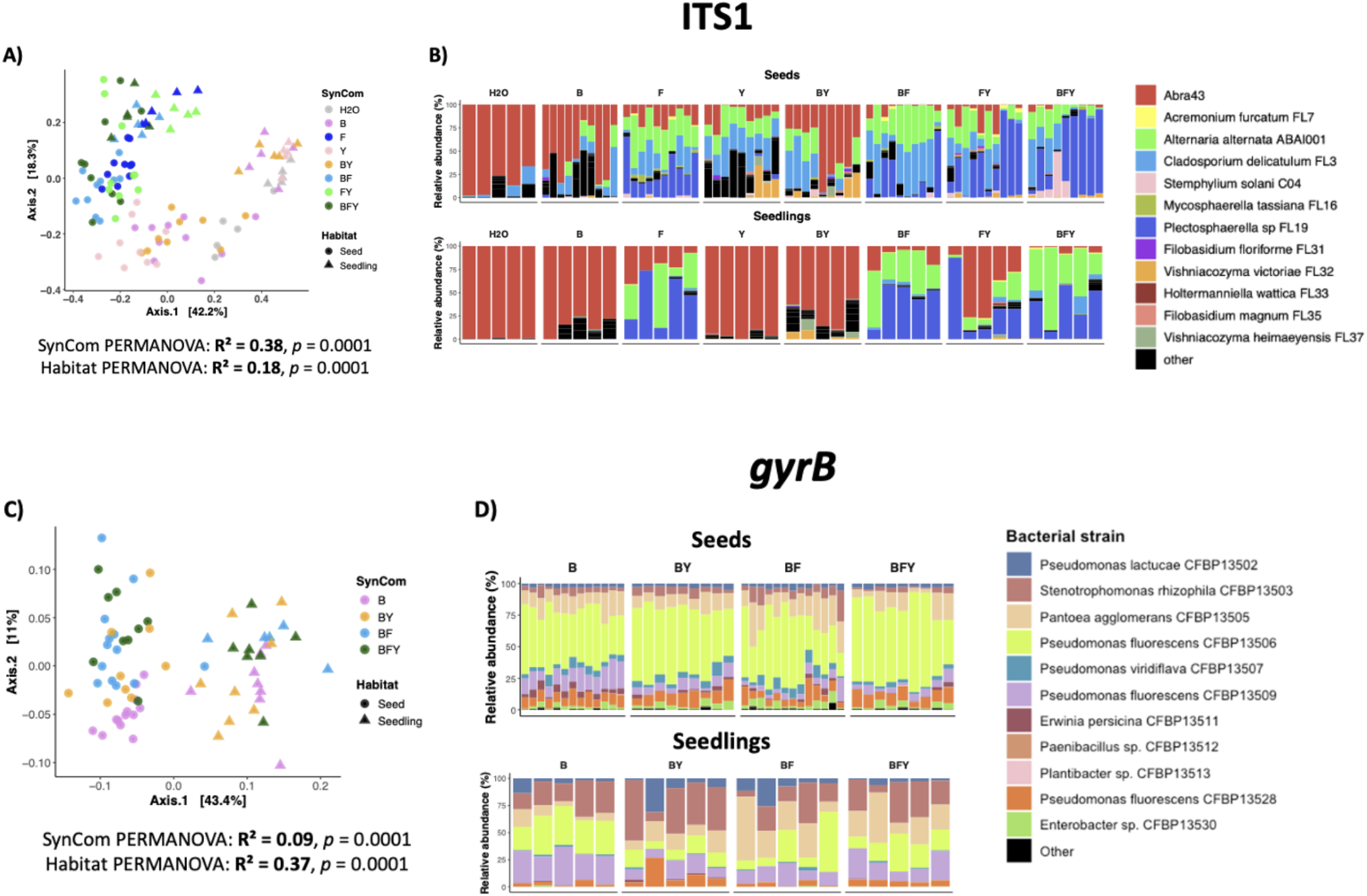
SynCom effects on fungal and bacterial community composition in seeds and seedlings inoculated with Abra43. A) Principal coordinate analysis based on Bray-Curtis dissimilarities of *ITS1* region fungal community profiles from individual seeds and seedlings. Points represent individual samples, colours indicate SynCom treatment, and shapes indicate habitat: seeds or seedlings. PERMANOVA showed significant effects of both SynCom treatment and habitat on fungal community composition (SynCom: R^2^ = 0.38, *p* = 0.0001; habitat: R^2^ = 0.18, *p* = 0.0001). B) Relative abundance profiles of fungal taxa in seeds and seedlings. Each stacked bar represents one individual sample. Colours indicate *Abra43*, fungal and yeast SynCom members, and taxa grouped as “other.” C) Principal coordinate analysis based on Bray-Curtis dissimilarities of *gyrB* gene bacterial community profiles from individual seeds and seedlings. Only SynCom treatments containing the bacterial fraction were included in the analysis (B, BY, BF, and BFY), as no amplification was obtained for the H2O, F, Y, and FY treatments. PERMANOVA showed significant effects of SynCom treatment and habitat on bacterial community composition (SynCom: R^2^ = 0.09, *p* = 0.0001; habitat: R^2^ = 0.37, *P* = 0.0001). D) Relative abundance profiles of bacterial SynCom strains in seeds and seedlings.

In seedlings treated with SynCom F, the reduction of Abra43 relative abundance coincided primarily with increased representation of *Alternaria alternata* ABAI001 and *Plectosphaerella* sp. FL19 (Fig. 2B). Addition of the bacterial fraction in SynCom BF was associated with a further reduction in Abra43 dominance and a broader representation of fungal SynCom members, including *Cladosporium delicatulum* FL3. These patterns suggest that the bacterial fraction may influence the establishment or relative representation of fungal SynCom members (Durán et al., 2018; Wolinska et al., 2021), although the underlying interactions cannot be resolved from relative-abundance data alone (Morton et al., 2019).

Inoculated yeast strains remained poorly represented in seedling communities in SynCom Y (0.68 ± 0.34 % of relative abundance). Interestingly, when yeasts were combined with bacteria in SynCom BY, yeast strains were more readily detected (8.76 ± 3.6 % of relative abundance), particularly *Vishniacozyma victoriae* gRaFL32 (4.01 ± 1.92 % of relative abundance) and *Vishniacozyma* sp. gRaFL37 (4.06 ± 1.36 %), together with a greater contribution of resident fungal taxa from native seed microbiota grouped as “Other.” This pattern suggests that yeast establishment may depend on community context, consistent with the strong microbial selection occurring during seed germination and emergence (Barret et al., 2015). Positive bacterial-fungal associations during germination have also been shown to enhance fungal establishment or plant development (Zhao et al., 2024). Moreover, suppressive seed microbiomes have been associated with a higher relative abundance of basidiomycetous yeasts, including strains from *Vishniacozyma, Filobasidium*, and *Papiliotrema* genera, together with bacterial taxa such as *Massilia* (Diakaki et al., 2025), suggesting that cross-kingdom associations may also contribute to disease suppression. Together, these observations raise the possibility that bacterial partners facilitate the persistence or establishment of yeasts during germination. Such facilitation could involve the formation of mixed-species biofilms that enhance microbial adhesion and protection against environmental stresses, as well as trophic interactions or metabolite exchange that increase nutrient availability. These mechanisms are well documented in bacterial-fungal associations and have been proposed to contribute to microbial establishment and persistence in plant-associated habitats (Frey-Klett et al., 2011; van Overbeek and Saikkonen, 2016; Deveau et al., 2018). However, such facilitation would require direct experimental validation. In SynCom FY, Abra43 contribution remained more variable and generally higher than in SynCom F alone, indicating that the addition of yeasts did not systematically enhance the pathogen-suppressive phenotype of the fungal fraction.

The complete SynCom BFY displayed the lowest overall Abra43 dominance and supported the simultaneous representation of several fungal members, particularly *A. alternata* ABAI001 and *Plectosphaerella* sp. FL19, together with lower abundances of *C. delicatulum* FL3 and *Acremonium furcatum* FL7. Thus, the strongest protective phenotype coincided with a fungal community in which pathogen dominance was reduced and several SynCom members were able to establish simultaneously. This observation supports the idea that protective activity may emerge from community context rather than from the performance of a single dominant antagonist.

Bacterial communities followed a markedly different assembly pattern. Although SynCom composition significantly affected bacterial β-diversity (PERMANOVA, R^2^ = 0.09, p = 0.0001), habitat explained a substantially larger proportion of the variation (R^2^ = 0.37, p = 0.0001; Fig. 2C). Seeds and seedlings consequently formed clearly separated groups in the PCoA, whereas B, BY, BF, and BFY treatments showed considerable overlap within each habitat. These results indicate that the seed-to-seedling transition acted as a stronger ecological filter on bacterial community assembly than the initial composition of the inoculated SynCom. This is consistent with previous findings obtained with bacterial SynComs on radish seeds and multi-kingdom SynComs in rapeseed (Simonin et al., 2023, Suteau et al., 2026).

Taxonomic profiles supported this interpretation (Fig. 2D). Seed bacterial communities were relatively similar among B, BY, BF, and BFY treatments and were largely dominated by *Pseudomonas fluorescens* CFBP13506. Following germination, bacterial communities became more heterogeneous, with a reduced contribution of *P. fluorescens* CFBP13506 in several seedlings and increased representation of other inoculated strains, including *Stenotrophomonas rhizophila* CFBP13503 and *Pantoea agglomerans* CFBP13505. Germination therefore appears to impose strong ecological filtering on bacterial SynCom members, such that abundance on the seed does not necessarily predict persistence in the emerging seedling, as also observed in previous studies of seed-to-seedling microbial transmission (Rochefort et al., 2021; Chesneau et al., 2022). In contrast, fungal communities retained a stronger SynCom-dependent signature, possibly reflecting greater persistence of particular inoculated fungal strains across the seed-to-seedling transition. Seed-transmitted fungi can remain abundant in juvenile plant microbiota (Johnston-Monje et al., 2021), although such transmission is strongly taxon-dependent, suggesting that the pattern observed here may reflect strain-specific ecological competence rather than a general property of fungi. Together, these results reveal distinct assembly trajectories for the two microbial kingdoms during early plant development.

Interestingly, *A. alternata* ABAI001 and *Plectosphaerella* sp. FL19, which were well represented in several pathogen-suppressive SynComs, also displayed strong antagonistic activity against Abra43 in dual-culture *in vitro* assays on 2% Malt agar medium, with 58.14% and 44.37% growth inhibition, respectively (Fig. 3). However, *in vitro* antagonism did not consistently predict strain performance *in planta. Stemphylium solani* C04, for example, displayed substantial antagonistic activity against Abra43 *in vitro* (37.18% inhibition) and was detected on inoculated seeds, but its relative abundance markedly decreased during the seed-to-seedling transition, suggesting poor persistence despite its antagonistic potential. Similar discrepancies between *in vitro* antagonism and disease suppression *in planta* have been reported previously (Whitaker and Bakker, 2019). This contrast suggests that antagonistic potential alone is insufficient to predict the contribution of individual strains to SynCom-mediated disease suppression. Their contribution is also likely to depend on their ability to establish and persist during the seed-to-seedling transition and to function within the surrounding microbial community (Simonin et al., 2023; Emmenegger et al., 2023; Zhou et al., 2024).

**Fig. 3.**
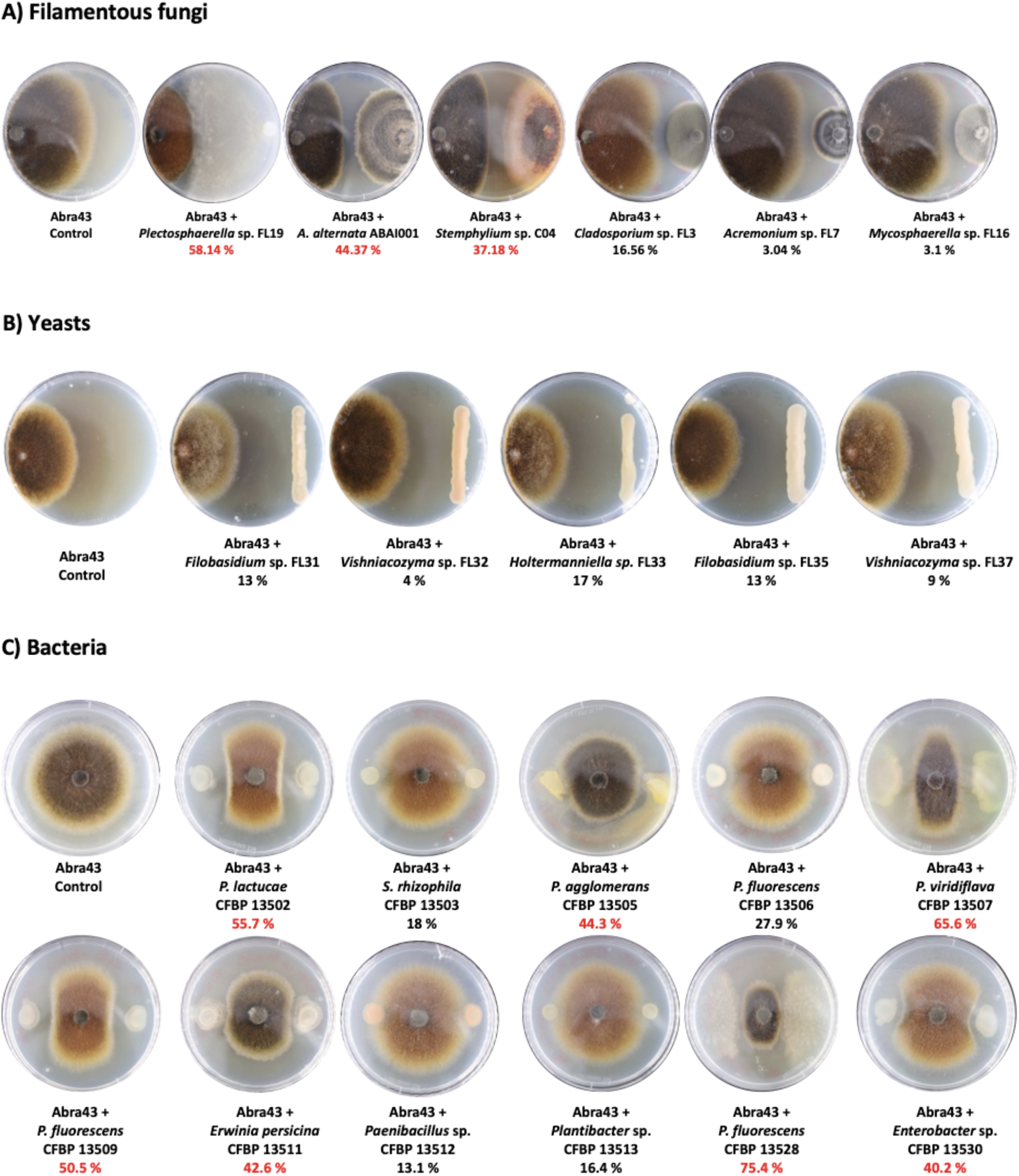
*in vitro* antagonistic activity of microorganisms-members of SynComs against Abra43. Representative dual-culture assays showing the growth of Abra43 alone (control) or in confrontation with (A) filamentous fungi, (B) yeasts, and (C) bacteria included in the synthetic communities (SynComs) (n = 3 replicates). Confrontation assays were performed on 2% malt agar. Abra43 was inoculated on one side of the plate and confronted with the corresponding microbial isolate.

Antagonistic activity was assessed by comparing Abra43 growth toward the interacting microorganism with its growth in the control. Numbers indicate Abra43 growth inhibition (%) relative to the control; values >35% are highlighted in red.

The contrasting effects of bacterial and fungal SynComs on seedling growth, symptom incidence, and pathogen abundance suggest that these microbial groups may contribute to plant protection through different mechanisms. Such functional complementarity across microbial kingdoms has been demonstrated in other plant SynCom systems, where bacterial and fungal communities contributed through both pathogen suppression and modulation of plant responses (Durán et al., 2018; Zhou et al., 2022). Seedlings treated with SynCom B maintained near-control growth despite substantial Abra43 colonization, a pattern consistent with a tolerance-like response rather than strong pathogen exclusion. Bacterial SynCom members may compensate for pathogen-induced growth inhibition by modulating phytohormone homeostasis and the growth-defense balance, thereby maintaining plant developmental processes despite infection (Finkel et al., 2020; Ma et al., 2021; Hou et al., 2021). Such an effect could additionally involve limitation of pathogen-induced physiological damage through maintenance of redox homeostasis and antioxidant capacity, as reported for beneficial bacteria in other *Alternaria*-plant interactions (Abdelhameed et al., 2024). Filamentous fungal-containing communities, by comparison, were more consistently associated with reduced pathogen relative abundance, suggesting a greater contribution to pathogen suppression. Combining these two microbial fractions may therefore broaden the range of protective functions available to the host. Consistent with this interpretation, the BF SynCom combined reduced Abra43 abundance with improved seedling growth. Similar functional complementarity has been reported in tomato, where bacterial-fungal SynComs provided stronger protection against Fusarium wilt than either bacterial or fungal communities alone (Zhou et al., 2022).

The contribution of the yeast fraction appears particularly context dependent. Yeasts alone showed no evidence of protection, whereas the addition of yeasts to the bacterial-fungal community resulted in the BFY SynCom, which combined reduced disease incidence, restored seedling growth, and the lowest pathogen colonization, making it the most protective treatment observed in this study. Thus, the absence of protection by SynCom Y alone does not exclude a contribution of yeasts when embedded within a more complex multi-kingdom community. Experimental reconstruction of plant microbiota has shown that microbial functions and establishment can depend strongly on interactions with members of other microbial kingdoms (Durán et al., 2018). In our system, yeast members may therefore become functionally relevant only in the presence of bacterial and fungal partners, for example through facilitation of their establishment, altered resource availability, or indirect modification of yeast-plant or yeast-pathogen interactions. However, our data do not demonstrate a direct beneficial interaction involving yeasts, and targeted experiments will be required to determine whether their contribution to BFY protection results from improved persistence or other cross-kingdom interactions.

Overall, our results suggest that protective seed-applied SynComs should combine functional potential with ecological competence rather than rely on community complexity alone. The bacterial fraction was associated primarily with maintenance of seedling growth despite substantial Abra43 colonization, whereas filamentous fungi contributed more directly to pathogen suppression, with *A. alternata* ABAI001 and *Plectosphaerella* sp. FL19 combining *in vitro* antagonism with successful establishment *in planta*. The superior performance of BFY further suggests that community context can modify the contribution of individual microbial groups, including yeasts that were ineffective when applied alone. Future SynCom design should therefore screen candidate strains along at least two complementary axes: antagonistic or plant-protective activity and the ability to establish and persist in the target habitat. Strain identity and community context should then be considered when assembling combinations, as these properties can be stronger predictors of protection than general community characteristics such as phylogenetic diversity or evenness (Emmenegger et al., 2023).

## Materials and Methods

Seeds of *Raphanus sativus* cv. Flamboyant 5 (Vilmorin-Mikado) were surface-disinfected by immersion in 70% ethanol for 1 min followed by sodium hypochlorite for 5 min, rinsed five times with sterile water, and dried for 15 min under a laminar-flow hood. The SynCom strains originated from the cultivable core microbiota of radish seeds. The bacterial fraction comprised 11 strains from the CIRM-CFBP collection: *Pseudomonas lactucae* CFBP13502, *Stenotrophomonas rhizophila* CFBP13503, *Pantoea agglomerans* CFBP13505, *Pseudomonas fluorescens* CFBP13506, *Pseudomonas viridiflava* CFBP13507, *Pseudomonas fluorescens* CFBP13509, *Erwinia persicina* CFBP13511, *Paenibacillus* sp. CFBP13512, *Plantibacter* sp. CFBP13513, *Pseudomonas fluorescens* CFBP13528 and *Enterobacter* sp. CFBP13530. The fungal and yeast fractions included 6 filamentous fungi: *Acremonium furcatum* FL7, *Alternaria alternata* ABAI001, *Cladosporium delicatulum* FL3, *Stemphylium* sp. C04, *Mycosphaerella* sp. FL16, *Plectosphaerella* sp FL19 and 5 yeasts: *Filobasidium* sp. gRaFL31, *Vishniacozyma victoriae* gRaFL32, *Holtermanniella takashimae* gRaFL33, *Filobasidium* sp. gRaFL35, *Vishniacozyma* sp. gRaFL37, were isolated in FungiSem lab and stored in our collection. Each bacterial strain was distinguishable by a specific *gyrB* ASV and each fungal or yeast strain by a specific ITS1 ASV, allowing their monitoring by metabarcoding. *Alternaria brassicicola* Abra43, originally isolated from radish seeds (Belmas et al., 2018), was used as the seed-borne pathogen.

Seven SynComs were assembled from three groups of microorganisms: bacteria (B), filamentous fungi (F), and yeasts (Y). These included each microbial group alone (B, F, and Y), all possible two-groups combinations (BF, BY, and FY), and the complete three-groups SynCom (BFY). Bacterial and yeast suspensions were adjusted to 10^6^ CFU/mL, and filamentous fungi and Abra43 suspensions to 10^6^ conidia/mL. Strains were mixed in equal proportions within each microbial fraction. For multi-kingdom SynComs, equal volumes of each microbial fraction were combined, and the final volume of all SynCom preparations was adjusted to the same volume with sterile water. Thus, the volume of inoculum applied to seeds was identical across treatments (10 mL for 100 seeds). For pathogen-inoculated treatments, Abra43 was added to the SynCom suspension, whereas an equivalent volume of sterile water was added to pathogen-free controls.

Surface-disinfected seeds (n = 100 per SynCom treatment) were immersed in the inocula for 30 min at 20°C, dried for 15 min, and incubated for 9 days at 20°C under a 16 h light/8 h dark photoperiod. All seedlings (n = 100 per SynCom treatment) were measured and scored for disease symptoms, and eight individual seedlings per treatment were sampled for microbiota analysis.

DNA was extracted using the NucleoSpin 96 Food Kit (Macherey-Nagel) following the manufacturer’s recommendations. Fungal and bacterial communities were characterized by ITS1 and *gyrB* gene amplicon sequencing, respectively, using an Illumina MiSeq platform, as previously described (Barret et al., 2015).

Sequences were processed with *cutadapt* and DADA2, fungal ASVs were assigned using the UNITE database, and bacterial ASVs using our curated *gyrB* gene database. Contaminants and low-abundance ASVs were removed before downstream analyses. Community composition was analyzed using Bray-Curtis dissimilarities followed by principal coordinate analysis (PCoA), and treatment effects were tested by PERMANOVA using the adonis2 function of the vegan package.

Abra43 abundance in seedlings was additionally quantified by qPCR targeting the pathogen-specific *dhn1* gene using primers *Ab*.*dhn1F-q* (ATGGGTCAGGTTGGCTGTAG) and *Ab*.*dhn1R-q* (ACGCAGATCACCCCAATATC). Differences among SynCom treatments were assessed using Kruskal-Wallis tests followed by Dunn’s multiple-comparison tests with Benjamini-Hochberg correction.

A more detailed Materials and Methods section is available in the “e-Xtra Supplementary Material”.

## Supporting information

e-Xtra Supplementary Material

## Acknowledgments

We thank Muriel Bahut (ANAN platform, SFR QuaSAV) for amplicon sequencing and International Centre of Microbial Resource (CIRM) - French Collection for Plant-associated Bacteria. INRAE. https://doi.org/10.15454/E8XX-4Z18

## Literature Cited

1. Barret, M., Briand, M., Bonneau, S., Préveaux, A., Valière, S., Bouchez, O., Hunault, G., Simoneau, P., and Jacques, M.-A. 2015. Emergence shapes the structure of the seed microbiota. Applied and Environmental Microbiology 81:1257–1266. DOI: 10.1128/AEM.03722-14.

2. Chesneau, G., Laroche, B., Préveaux, A., Marais, C., Briand, M., Marolleau, B., Simonin, M., and Barret, M. 2022. Single Seed Microbiota: Assembly and Transmission from Parent Plant to Seedling. mBio 13:e01648–22. DOI: 10.1128/mbio.01648-22.

3. Chirco, E. M., and Harman, G. E. 1978. The effects of Alternaria brassicicola infection on Brassica seed vigor and viability. J. Seed Technol. 3:12–22.

4. De Britto, S., and Jogaiah, S. 2022. Priming with fungal elicitor elicits early signaling defense against leaf spot of broccoli underlying cellular, biochemical and gene expression. Microbiological Research 263:127143. DOI: 10.1016/j.micres.2022.127143.

5. Durán, P., Thiergart, T., Garrido-Oter, R., Agler, M., Kemen, E., Schulze-Lefert, P., and Hacquard, S. 2018. Microbial interkingdom interactions in roots promote Arabidopsis survival. Cell 175:973–983.e14. DOI: 10.1016/j.cell.2018.10.020.

6. Emmenegger, B., Massoni, J., Pestalozzi, C. M., Bortfeld-Miller, M., Maier, B. A., et al. 2023. Identifying microbiota community patterns important for plant protection using synthetic communities and machine learning. Nature Communications 14:7983.

7. Guillemette, T., Calmes, B., and Simoneau, P. 2014. Impact of the UPR on the virulence of the plant fungal pathogen A. brassicicola. Virulence 5:357–364. DOI: 10.4161/viru.26772.

8. Iacomi-Vasilescu, B., Avenot, H., Bataille-Simoneau, N., Laurent, E., Guénard, M., and Simoneau, P. 2004. In vitro fungicide sensitivity of Alternaria species pathogenic to crucifers and identification of Alternaria brassicicola field isolates highly resistant to both dicarboximides and phenylpyrroles. Crop Protection 23:481–488. DOI: 10.1016/j.cropro.2003.10.003.

9. Kaur, N., and Dutta, B. 2024a. Aggressive Alternaria brassicicola with reduced fungicide sensitivity can be associated with naturally infested broccoli seeds. Plant Disease 108:2154–2161. DOI: 10.1094/PDIS-01-24-0056-RE.

10. Kaur, N., and Dutta, B. 2024b. Characterization of seed-to-seedling transmission of Alternaria brassicicola in broccoli. Plant Disease 108:2046–2052. DOI: 10.1094/PDIS-10-23-2002-RE.

11. Liu, Q., Cheng, L., Zhang, E., Ling, L., Tan, W., Liang, S., Shu, C., Ma, Q., Zhao, S., Wei, J., Wang, Y., Nian, H., Cheng, Y., and Lian, T. 2026. Cross-niche metabolite-microbiome interactions orchestrate systemic soybean resistance to Fusarium root rot. The ISME Journal 20(1):wrag080. DOI: 10.1093/ismejo/wrag080.

12. Marchi, M., Bosc-Bierne, A., Lerenard, T., Le Corff, J., Aligon, S., Rolland, A., Simonin, M., Marais, C., Briand, M., Cordovez, V., Gouka, L., Guillemette, T., Simoneau, P., and Guschinskaya, N. 2024. Unexplored yeast diversity in seed microbiota. bioRxiv. 2024.11.27.625647. DOI: 10.1101/2024.11.27.625647.

13. Matsumoto, H., Fan, X., Wang, Y., Kusstatscher, P., Duan, J., Wu, S., Chen, S., Qiao, K., Wang, Y., Ma, B., Zhu, G., Hashidoko, Y., Berg, G., Cernava, T., and Wang, M. 2021. Bacterial seed endophyte shapes disease resistance in rice. Nature Plants 7:60–72. DOI: 10.1038/s41477-020-00826-5.

14. Pattanamahakul, P., and Strange, R. N. 1999. Identification and toxicity of Alternaria brassicicola, the causal agent of dark leaf spot disease of Brassica species grown in Thailand. Plant Pathology 48:749–755. DOI: 10.1046/j.1365-3059.1999.00406.x.

15. Pochon, S., Terrasson, E., Guillemette, T., Iacomi-Vasilescu, B., Georgeault, S., Juchaux, M., Berruyer, R., Debeaujon, I., Simoneau, P., and Campion, C. 2012. The Arabidopsis thaliana-Alternaria brassicicola pathosystem: A model interaction for investigating seed transmission of necrotrophic fungi. Plant Methods 8:16. DOI: 10.1186/1746-4811-8-16.

16. Rochefort, A., Simonin, M., Marais, C., Guillerm-Erckelboudt, A.-Y., Barret, M., and Sarniguet, A. 2021. Transmission of Seed and Soil Microbiota to Seedling. mSystems 6:e00446–21. DOI: 10.1128/mSystems.00446-21.

17. Rybakova, D., Mancinelli, R., Wikström, M., Birch-Jensen, A.-S., Postma, J., Ehlers, R.-U., Goertz, S., and Berg, G. 2017. The structure of the Brassica napus seed microbiome is cultivar-dependent and affects the interactions of symbionts and pathogens. Microbiome 5:104. DOI: 10.1186/s40168-017-0310-6.

18. Saharan, G. S., Mehta, N., and Meena, P. D. 2016. Alternaria Diseases of Crucifers: Biology, Ecology and Disease Management. Springer, Singapore. DOI: 10.1007/978-981-10-0021-8.

19. Simonin, M., Préveaux, A., Marais, C., Garin, T., Arnault, G., Sarniguet, A., and Barret, M. 2023. Transmission of synthetic seed bacterial communities to radish seedlings: impact on microbiota assembly and plant phenotype. Peer Community Journal 3:e95. DOI: 10.24072/pcjournal.329.

20. Stincone, P., Braun, L. M., Bağcı, C., Navarro-Diaz, M., Pérez-Lorente, A. I., Farrell, S. P., Gómez-Pérez, D., Bode, J., Steuer-Lodd, K., Mahmoudi, M., Chaudhry, V., Romero, D., Aron, A. T., Ziemert, N., Molina-Santiago, C., Kemen, E. M., and Petras, D. 2026. Cooperative siderophore use stabilizes a protective leaf microbiome. bioRxiv [Preprint]. 2026.03.18.712463. DOI: 10.64898/2026.03.18.712463.

21. Suteau, L., Colaert-Sentenac, L., Law, S. R., and Simonin, M. 2025. Transient microbial architects: tracing the legacy effects of ephemeral taxa during plant microbiome assembly. Current Opinion in Microbiology 88:102664. DOI: 10.1016/j.mib.2025.102664.

22. Suteau, L., Campion, C., Marais, C., Briand, M., Hardouin, A., Hellyn, K., Maurice, K., Marchi, M., Simonin, M., and Guschinkaya, N. 2026. Seed-applied multi-kingdom synthetic communities selectively reshape bacterial communities and highlight key criteria for strain selection. bioRxiv. 2026.07.15.738658. DOI: 10.64898/2026.07.15.738658.

23. Walitang, D. I., Kim, K., Madhaiyan, M., Kim, Y. K., Kang, Y., and Sa, T. 2017. Characterizing endophytic competence and plant growth promotion of bacterial endophytes inhabiting the seed endosphere of rice. BMC Microbiology 17:209. DOI: 10.1186/s12866-017-1117-0.

24. Zhou, X., Wang, J., Liu, F., Liang, J., Zhao, P., Tsui, C. K. M., et al. 2022. Cross-kingdom synthetic microbiota supports tomato suppression of Fusarium wilt disease. Nature Communications 13:7890.

