## Supplementary material for "A Seed-Derived Multi-Kingdom Synthetic Community Enhances Radish Seedling Protection Against *Alternaria brassicicola* Abra43": e-Xtra Supplementary Material

### Plant material

Seeds used for this experiment were of the radish genotype *Raphanus sativus* "Flamboyant 5" (Vilmorin Mikado). Prior to the experiment, approximately the seeds were surface-disinfected: first immersed in 70% ethanol for one minute, followed by a 5-minute soaking in sodium hypochlorite, and then rinsed with sterile water 5 times. The seeds were subsequently dried in a laminar flow hood for 15 minutes.

### Microbial strains of SynComs

#### Bacterial strains

All bacterial strains used in this study were supplied by the French Collection of Plant-Associated Bacteria (CIRM-CFBP). Eleven strains were selected based on their previously identified status as members of the core microbiota of radish seeds (Simonin et al., 2022). The selected bacterial set included: *Pseudomonas lactucae* CFBP13502, *Stenotrophomonas rhizophila* CFBP13503, *Pantoea agglomerans* CFBP13505, *Pseudomonas fluorescens* CFBP13506, *Pseudomonas viridiflava* CFBP13507, *Pseudomonas fluorescens* CFBP13509, *Erwinia persicina* CFBP13511, *Paenibacillus* sp. CFBP13512, *Plantibacter* sp. CFBP13513, *Pseudomonas fluorescens* CFBP13528 and *Enterobacter* sp. CFBP13530 (Figures S1). Each bacterial strain possessed a unique *gyrB* ASV, enabling strain-level monitoring by metabarcoding. The strains were routinely cultured and maintained on 10% Tryptic Soy agar (TSA) at 20 °C. Antagonistic activity against *A. brassicicola* Abra43 was first assessed *in vitro* on 2% Malt extract agar.

### Filamentous fungi and yeast strains

The *A. brassicicola* strain used was the Abra43 strain which was isolated in 1999 on a radish seed during quality controls of French commercial lots and purified by monospore isolation and identified as *A. brassicicola* based on morphological and molecular analyses (Avenot et al. 2005; Belmas et al. 2018). This strain exhibits a strong aggressiveness on cabbage and *Arabidopsis* sp. leaves and is also efficiently transmitted to seeds (Stephanie Pochon et al. 2012) (Pochon et al., 2012; Rezki et al., 2016). As the other strains, Abra43 was cultivated and maintained on malt agar (MA) at 20°C. Antagonistic activity against *A. brassicicola* Abra43 was first assessed *in vitro* on 2% malt extract agar.

SynCom filamentous fungi and yeast strains were isolated from *R. sativus* seeds by our team. When all strains were pure, they were all identified with the ITS1 genetic marker. DNA was extracted from the strains with the NucleoSpin® 96 DNA from the food de Macherey-Nagel kit proceeding as instructed by the given protocol. ITS1 region was then amplified by PCR with the use of ITS4 and NS7 primers (Simonin et al. 2026). First the DNA was denatured for 3 minutes at 96°C, then 35 cycles of 30 seconds of denaturation at 96°C, 30 seconds of primer linking at 56°C and 2 minutes 30 seconds of extensions at 72°C were performed before concluding by 5 minutes of extension still at 72°C. Samples were sent for Sanger sequencing (Eurofins). Results received were sorted and sequences with low quality were removed. Sequences were cut in order to only keep the region between the ITS1-F and ITS2-R primers to match to ITS1 sequences used in metabarcoding to monitor the strains in the seed and seedling compartments. The sequences were taxonomically assigned using the R package dada2 using the UNITE V8.3 database as reference. Taxonomic assignment was then used to select 6 fungi strains (all sporulating) and 5 yeast strains all belonging to the core microbiota of *R. sativus* seeds (Table S1). Strains were then cultivated and maintained on 2% Malt extract agar at 20°C. Each fungal strain possessed a unique ITS1 ASV, enabling strain-level monitoring by metabarcoding. Antagonistic activity against *A. brassicicola* Abra43 was first assessed *in vitro* on 2% malt extract agar. Culture filtrates from each bacterial strain were subsequently tested against Abra43 using nephelometric measurements. Finally, the protective capacity of each strain was evaluated by co-inoculation on radish seeds to determine whether it could reduce disease development caused by the fungal pathogen.

**SynComs preparation and inoculation**

Seven SynComs of increasing complexity were prepared: three single-kingdom communities containing bacteria (B), filamentous fungi (F), or yeasts (Y); three two-kingdom communities (BF, BY, and FY); and one three-kingdom community (BFY). Bacterial cells were harvested from 10% TSA plates, resuspended in sterile water, and adjusted to $10^6$ CFU/ml on OD600 measurements. Yeast cells were harvested from 2% malt agar plates, resuspended in sterile water, quantified using a Thoma counting chamber, and adjusted to $10^6$ CFU/ml. The microbial strains were then mixed in equal proportions to assemble the different SynComs. Conidial suspensions of the filamentous fungi and Abra43 were prepared in sterile water, quantified using a Thoma counting chamber, and adjusted to $10^6$ conidia/ml.

SynCom inocula were plated on 2% malt agar and 10% TSA to confirm the viability of their microbial members. In addition, 200 µL of each inoculum was stored at −20 °C for subsequent metabarcoding analysis. Each SynCom was divided into two experimental conditions. The Abra43 conidial suspension was added to the pathogen-inoculated condition, whereas an equivalent volume of sterile water was added to the pathogen-free condition. Surface-disinfected radish seeds were immersed in the inoculum and agitated for 30 min at 20 °C, then dried for 15 min. Approximately 50 seeds per treatment were retained for seed microbiota analysis, while the remaining 100 seeds were placed in sterile pleated-paper germination boxes and incubated for 9 days at 20 °C under a 16 h light/8 h dark photoperiod.

**Seed and seedling sample collection**

The microbiota of individual seeds and seedlings was analyzed in response to the different SynCom treatments and pathogen inoculation. For seed samples, 10 individual seeds per condition were placed separately in the wells of a nephelometer plate. Each seed was immersed in 250 µL of sterile water and macerated by agitation for 2.5 h at 4 °C. After maceration, 200 µL of each suspension was collected and stored at −20 °C until metabarcoding analysis.

At 9 days post-inoculation, seedlings were photographed, measured, and assessed for symptoms caused by Abra43. For each condition, five individual seedlings were collected and placed separately in stomacher bags. After the addition of 500 µL of sterile water, the bags were sealed and the seedlings were homogenized. Finally, 200 µL of each homogenate was collected and stored at −20 °C until metabarcoding analysis.

**Amplicon library preparation and sequencing**

DNA from all collected samples was extracted using the NucleoSpin® 96 Food Kit from Macherey-Nagel® following the provided protocol. Genetic markers ITS1 and *gyrB* were then amplified by PCR using primers with MiSeq adapter sequences. For this PCR, the primers were used at a concentration of 10 µM with AccuPrime™ Taq DNA Polymerase High Fidelity, REF 12346-086, from Invitrogen by Thermo Fisher®. The PCR for both markers consisted of five stages: an initial denaturation at 94°C for 3 minutes, followed by 35 cycles of 30 seconds at 94°C for denaturation, 45 seconds at 50°C for primer annealing, and 1 minute 30 seconds at 68°C for extension, ending with a final extension period of 10 minutes at 68°C. Amplification of each sample was verified by gel electrophoresis before the samples were purified using Sera-Mag beads with a DNA bead solution ratio of 1:1. A second PCR was then conducted to integrate indexing sequences for sample identification. The primers (see Appendix 1) for this PCR were also prepared at 10 µM, using the same Taq polymerase as in the previous PCR. This PCR consisted of 5 stages: 60 seconds of initial denaturation at 94°C, followed by 12 cycles of

60 seconds at 94°C for denaturation, 60 seconds at 55°C for primer annealing, and 60 seconds at 68°C for extension, ending with a final extension period of 10 minutes at 68°C. Amplification of some samples was verified by gel electrophoresis, and samples were purified using Sera-Mag beads with a DNA bead solution ratio of 1:1. Some samples were then quantified using the Quanti™ PicoGreen™ dsDNA Assay Kit. Following this quantification, it was decided to add twice as much of the 16S amplicons to the equimolar pool as the ITS1 amplicons. The purity of the pool was verified on a gel before it was diluted to one-fifth and quantified with the KAPA Library Quant Kit for Illumina on the eve of sequencing. The sequencing of the pool was performed on the ANAN platform's on an Illumina MiSeq device at the SFR QuaSAV in Angers. A MiSeq 600V3 cartridge was used for the run, and 14pM was loaded onto the cartridge.

**Data analysis from sequencing run**

Bioinformatic processing of the amplicons was performed in R (v4.5.2). Primer sequences were removed using cutadapt v2.7 (Martin 2011) and trimmed FASTQ files were processed with DADA2 v1.22.0 (Callahan et al. 2016) forward and reverse sequences were then assembled and underwent taxonomic assignment. Amplicon Sequence Variant (ASV) were taxonomically assigned using a naïve Bayesian classifier (Wang et al. 2007) with an in-house *gyrB* database (train_set_gyrB_v5.fa.gz, available upon request). Unassigned sequences at the phylum level and *parE* sequences (a *gyrB* paralog) were removed. ITS1-derived ASVs were classified against the UNITE 8.3 database (Abarenkov et al. 2024) and non-fungal ASVs were filtered out. All scripts are available at XXX. Then, post-clustering curation was performed using LULU v0.1.0 (Frøslev et al. 2017) at a minimum threshold of 98% for *gyrB* and 99% for ITS. Contaminants were identified and removed using the decontam package v1.30.0, followed by manual verification and cleaning by analyzing positive and negative sequencing controls. ASVs with fewer than 10 reads and detected in only one sample were removed from the dataset. Finally, samples with fewer than 1000 reads after filtering were excluded from downstream analysis. To assess sequencing completeness, rarefaction curves were generated using the rarecurve function of the vegan package (v2.7.2 10.32614/CRAN.package.vegan) (Figure S2). Based on rarefaction curves, the *gyrB* dataset was rarefied at 3000 reads, and ITS dataset was rarefied at 2600 reads. Rarefied ASV tables were then used to estimate species richness. For all other analyses, non-rarefied read counts were transformed into proportions (relative abundance) using the transform function from the microbiome R package (Lahti and Shetty 2017). To test the effect of SynCom inoculation on the microbiota, ordination analyses were performed using Bray-Curtis dissimilarity matrices computed on log(x+1) transformed non-rarefied data followed by Principal Coordinate Analysis (PCoA) using the ordinate function in phyloseq. To explore the percentage of variance explained by the factor SynCom on seed samples and seedling sample separately, permutational multivariate analysis of variance (PERMANOVA) was performed with 999 permutations using the adonis2 function from the vegan R package (Oksanen et al. 2012).

**Evaluation of the presence of *A. brassicicola* Abra43 on infected seedlings by qPCR**

To assess the influence of SynComs on the amount of *A. brassicicola* Abra43 present on the infected seedlings, we performed absolute quantitative PCR analyses (Brankatschk et al. 2012). The concentration of *A. brassicicola* Abra43 DNA used in the standard curve was assessed using the Invitrogen$^{TM}$ Qubit$^{TM}$ 4 Fluorometer (ANAN platform). For each SynCom condition, the DNA extracted from 5 seedlings was used to perform the quantification. All assays were performed on the thermocycler StepOnePlus Real Time PCR system (Applied Biosystem$^{TM}$) using the protocol described in Sellam et al. (2006). The primer pair Ab.dhn1F-q (5'-ACGCAGATCACCCCAATATC-3') and Ab.dhn1R-q (5'-ATGGGTCAGGTTGGCTGTAG) (Pochon et al. 2013) was used to quantify the presence of *A. brassicicola* on the seedlings ($R^2$ = 0,9998 and Amplification Factor = 1,85). The specificity of the *DHN1* primers on

*A. brassicicola* Abra43 was confirmed using the healthy SynCom samples, no amplification was detected.

The results in gene copy numbers of *A. brassicicola* were calculated as followed:

$$Gene\ copy\ number\ = \frac{(DNA\ concentration\ (ng/\mu L) \times Avogadro's\ number)}{A.\ brassicicola\ genome\ size\ (bp) \times conversion\ factor\ to\ ng \times average\ weight\ of\ a\ base\ par\ (Da)}$$

$$Gene\ copy\ number\ = \frac{(DNA\ concentration\ (ng/\mu L)\ \times\ [6.022\ \times\ 10^{23}]}{31036461\ \times [1 \times 10^{9}]\ \times\ 650}$$

Those results were compared between the SynCom conditions using the non-parametric test Kruskal-Wallis and Dunn’s test of multiple comparisons with Benjamini-Hochberg adjustment.


Abarenkov, Kessy, R. Henrik Nilsson, Karl-Henrik Larsson, et al. 2024. ‘The UNITE Database for Molecular Identification and Taxonomic Communication of Fungi and Other Eukaryotes: Sequences, Taxa and Classifications Reconsidered’. *Nucleic Acids Research* 52 (D1): D791–97. https://doi.org/10.1093/nar/gkad1039.

Avenot, Hervé, Anita Dongo, Nelly Bataillé-Simoneau, et al. 2005. ‘Isolation of 12 Polymorphic Microsatellite Loci in the Phytopathogenic Fungus Alternaria Brassicicola’. *Molecular Ecology Notes* 5 (4): 948–50. https://doi.org/10.1111/j.1471-8286.2005.01125.x.

Belmas, Elodie, Martial Briand, Anthony Kwasiborski, et al. 2018. ‘Genome Sequence of the Necrotrophic Plant Pathogen Alternaria Brassicicola Abra43’. *Genome Announcements* 6 (6): e01559-17. https://doi.org/10.1128/genomeA.01559-17.

Brankatschk, Robert, Natacha Bodenhausen, Josef Zeyer, and Helmut Bürgmann. 2012. ‘Simple Absolute Quantification Method Correcting for Quantitative PCR Efficiency Variations for Microbial Community Samples’. *Applied and Environmental Microbiology* 78 (12): 4481–89. https://doi.org/10.1128/AEM.07878-11.

Callahan, Benjamin J., Paul J. McMurdie, Michael J. Rosen, Andrew W. Han, Amy Jo A. Johnson, and Susan P. Holmes. 2016. ‘DADA2: High-Resolution Sample Inference from Illumina Amplicon Data’. *Nature Methods* 13 (7): 581–83. https://doi.org/10.1038/nmeth.3869.

Frøslev, Tobias Guldberg, Rasmus Kjøller, Hans Henrik Bruun, et al. 2017. ‘Algorithm for Post-Clustering Curation of DNA Amplicon Data Yields Reliable Biodiversity Estimates’. *Nature Communications* 8 (1): 1188. https://doi.org/10.1038/s41467-017-01312-x.

Lahti, Leo, and Sudarshan Shetty. 2017. *Microbiome R Package*. https://doi.org/10.18129/B9.bioc.microbiome.

Martin, Marcel. 2011. 'Cutadapt Removes Adapter Sequences from High-Throughput Sequencing Reads'. *EMBnet.Journal* 17 (1): 10–12. https://doi.org/10.14806/ej.17.1.200.

Oksanen, Jari, F. Guillaume Blanchet, Roeland Kindt, et al., dirs. 2012. *Vegan: Community Ecology Package*.

Pochon, Stéphanie, Philippe Simoneau, Sandrine Pigné, et al. 2013. 'Dehydrin-like Proteins in the Necrotrophic Fungus Alternaria Brassicicola Have a Role in Plant Pathogenesis and Stress Response'. *PLOS ONE* 8 (10): e75143. https://doi.org/10.1371/journal.pone.0075143.

Pochon, Stephanie, Emmanuel Terrasson, Thomas Guillemette, et al. 2012. 'The Arabidopsis Thaliana-Alternaria Brassicicola Pathosystem: A Model Interaction for Investigating Seed Transmission of Necrotrophic Fungi'. *Plant Methods* 8 (1): 16. https://doi.org/10.1186/1746-4811-8-16.

Sellam, Adnane, Pascal Poupard, and Philippe Simoneau. 2006. 'Molecular Cloning of AbGst1 Encoding a Glutathione Transferase Differentially Expressed during Exposure of Alternaria Brassicicola to Isothiocyanates'. *FEMS Microbiology Letters* 258 (2): 241–49. https://doi.org/10.1111/j.1574-6968.2006.00223.x.

Simonin, Marie, Natalia Guschinskaya, Muriel Marchi, et al. 2026. 'Seed Microbiota Diversity and Culture Collection of Four Major Crops Covering Different Genotypes and Production Modes'. Preprint, bioRxiv, May 1. https://doi.org/10.64898/2026.04.29.721552.

Wang, Qiong, George M. Garrity, James M. Tiedje, and James R. Cole. 2007. 'Naive Bayesian Classifier for Rapid Assignment of rRNA Sequences into the New Bacterial Taxonomy'. *Applied and Environmental Microbiology* 73 (16): 5261–67. https://doi.org/10.1128/AEM.00062-07.
